# Alpha-ketoglutarate, an endogenous metabolite, extends lifespan and compresses morbidity in aging mice

**DOI:** 10.1101/779157

**Authors:** Azar Asadi Shahmirzadi, Daniel Edgar, Chen-Yu Liao, Yueh-Mei Hsu, Mark Lucanic, Arash Asadi Shahmirzadi, Christopher Wiley, Rebeccah Riley, Brian Kaplowitz, Garbo Gan, Chisaka Kuehnemann, Dipa Bhaumik, Judith Campisi, Brian K Kennedy, Gordon J. Lithgow

## Abstract

The decline in early life mortality since the 1950s has resulted in dramatic demographic shift towards aged population. Aging manifests as a decline in health, multiple organ dysfunction and increased vulnerability to diseases, which degrades quality of life. A verity of genetic and pharmacological interventions, mostly from non-vertebrate models, have been identified that can enhance lifespan. Whether these interventions extend healthspan, the disease free and functional period of life, has only sometimes been tested and is often a matter of debate. Human aging indices have been developed to assess elements of functional decline with aging (e.g. sarcopenia, cognitive function). However, corresponding comprehensive indices in mice are seldom applied to aging studies. To probe the relationship between healthspan and lifespan extension in mammals, we performed a series of longitudinal, clinically-relevant healthspan measurements. Metabolism and aging are tightly connected and specific perturbations of nutrient-sensing pathways can enhance longevity in laboratory animals. Here we show that alpha-ketoglutarate (delivered in the form of a Calcium salt, CaAKG), a key metabolite in tricarboxylic (TCA) cycle that is reported to extend lifespan in worms, can significantly extend lifespan and healthspan in mice. AKG is involved in various fundamental processes including collagen synthesis and epigenetic changes. Due to its broad roles in multiple biological processes, AKG has been a subject of interest for researchers in various fields. AKG also influences several age-related processes, including stem cell proliferation and osteoporosis. To determine its role in mammalian aging, we administered CaAKG in 18 months old mice and determined its effect on the onset of frailty and survival, discovering that the metabolite promotes longer, healthier life associated with a decrease in levels of inflammatory factors. Interestingly the reduction in frailty was more dramatic than the increase in lifespan, leading us to propose that CaAKG compresses morbidity.

## Introduction

The decline in early life mortality since the 1950s has resulted in dramatic demographic shift towards an aged population. Aging manifests as a decline in health, multiple organ dysfunction and increased vulnerability to diseases, all of which degrade quality of life. A verity of genetic and pharmacological interventions, developed mostly from non-vertebrate model organisms, have been identified that enhance lifespan [1-4]. Whether these interventions extend healthspan, the disease free and functional period of life, has only sometimes been tested and is often a matter of debate [5-7]. Human aging indices have been developed to assess elements of functional decline with aging (e.g. sarcopenia, cognitive function). However, corresponding comprehensive indices in mice are seldom applied to aging studies [8]. To probe the relationship between healthspan and lifespan extension in mammals, we performed a series of longitudinal, clinically-relevant healthspan measurements.

Metabolism and aging are tightly connected and specific perturbations of nutrient-sensing pathways can enhance longevity in laboratory animals [9-11]. Here we show that alpha-ketoglutarate (delivered in the form of a calcium salt, CaAKG), a key metabolite in tricarboxylic acid (TCA) cycle that is reported to extend lifespan in worms [12], can significantly extend lifespan and healthspan in mice. AKG is involved in various fundamental processes, including central metabolism, collagen synthesis [13] and epigenetic regulation [14, 15]. Due to its broad roles in multiple biological processes, AKG has been a subject of interest for researchers in various fields [16]. AKG also influences several age-related processes, including stem cell proliferation [17, 18] and osteoporosis [19]. To study its role in mammalian aging, animals received sustained CaAKG treatment starting at 18 months of age and determined its effect on survival, as well as the onset of frailty. We find that CaAKG promotes longer, healthier life associated with a decrease in levels of inflammatory factors. Strikingly, the reduction in frailty was more dramatic than the increase in lifespan, leading us to propose that CaAKG compresses period of morbidity.

## Results

C57BL/6 mice were fed regular chow until they were started on a diet containing CaAKG at 540 days of life (Extended Diagram 1). We assessed two cohorts of mice each consisting of 45±2 females and 45±2 males (total of 182 animals), allowing us to check for reproducibility, one of the challenges in all areas of biology [20]. Here we report dietary supplemented 2% CaAKG (w/w) increases survival in two independent cohorts of aged mice (Fig. 1). In the first cohort of female mice, median lifespan and survival (age at 90^th^ percentile mortality) were significantly extended by 16.6% and 19.7% from inception of CaAKG feeding. These findings have been repeated in the second cohort of mice: the female median lifespan and survival were significantly extended by 10.5% and 8% (Fig. 1a, b and Extended Table 1). Although, improved survival for males was not significant in both cohorts (Fig. 1d, e), median lifespan was extended for 9.6%and 12.8% from inception of treatment, respectively (Extended Table 1).

**Figure 1.**
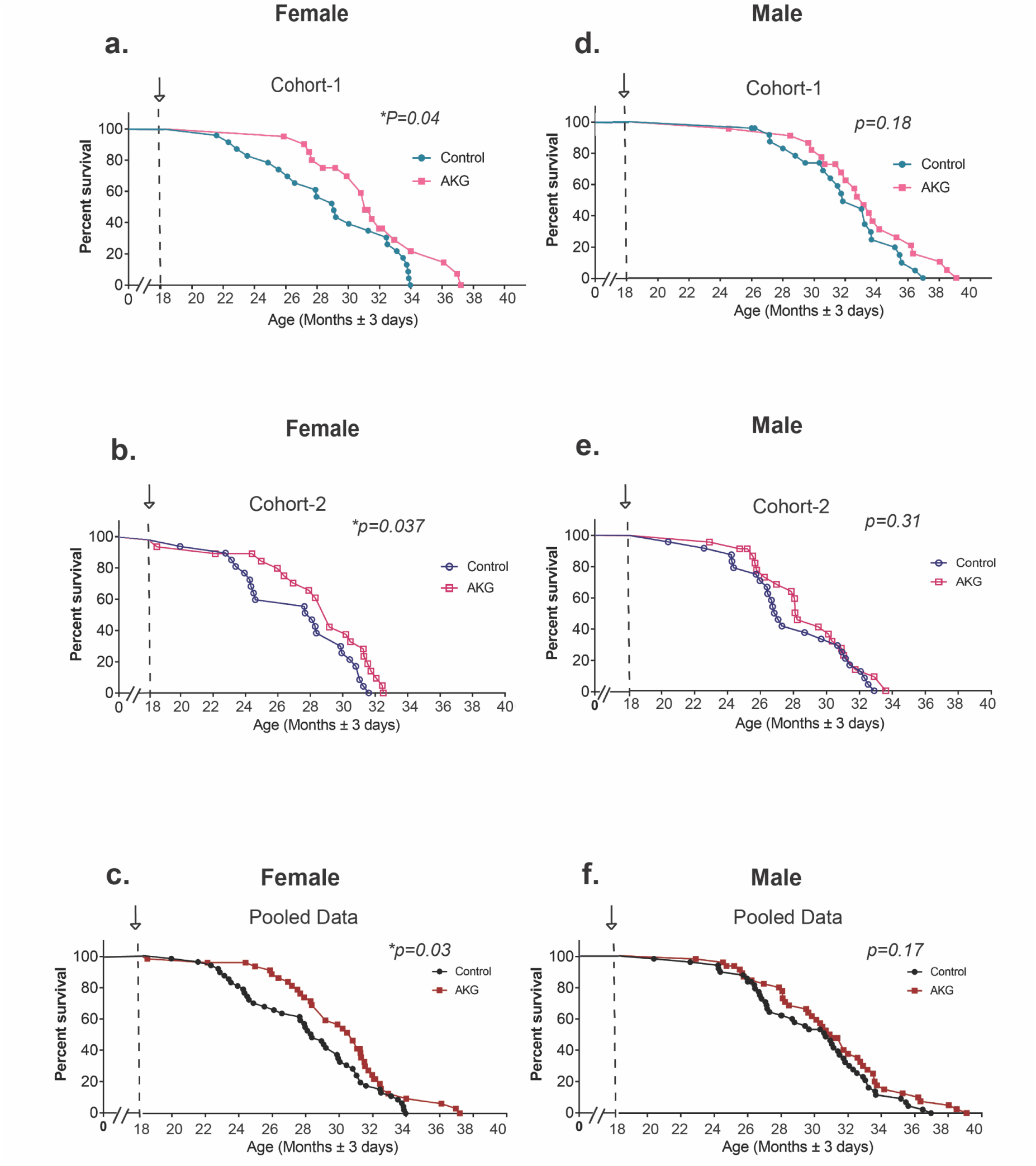
AKG extends lifespan and decrease mortality. Post treatment survival plots graphed for Cohort-1 and Cohort-2. Comparing control mice to those fed AKG in the diet starting at 18th months of age. Arrows indicate the start of the treatment. Survival curves for (**a**) Cohort-1 female (n=43), (**b**) Cohort-2 female (n=45), (**c**) Pooled female, (**d**) Cohort-1 male (n=46), (**e**) Cohort-2 male (n=47) and (**f**) Pooled male. Survival curve comparison were performed using Log-rank test, *P < 0.05. Maximum lifespan extensions were calculated using Fisher’s exact test statistics, **P =0.0064 for female Cohort-1 and *<0.021 for female Cohort-2.

In order to assess healthspan, we applied measurements that are based on a clinically relevant frailty index [8, 21]. The frailty index (FI) consists of 31 phenotypes that are indicators of age-associated health deterioration, with each phenotype scored on a 0, 0.5 or 1 scale, based on its severity. Body weight and surface temperature were collected and converted to the same scoring scale (Extended Table 1). All scorings were conducted in a blinded manner. The 31-metrics share many characteristics of the human frailty indices and have been reported to progress similarly with aging in mice and humans [22]. Measurements were repeated approximately every eight weeks, providing us with eight and seven sets of data respectively for male and female groups. To establish a baseline clinical assessment, we collected our dataset right before the start of the treatment at 18^th^ months (Fig. 2a-d). The total frailty score, which is the sum of 31 frailty phenotypes, indicates the state of increased vulnerability to adverse health outcomes, which is comparable to morbidity. In both female and male animals, CaAKG decreases incidence and severity of different aging phenotypes and postpones frailty (Fig. 2a, b). Females show significant health benefits after 9 months of CaAKG supplementation (P<0.001, Fig. 2a). After 11 months of treatment, male animals start to show significant health improvement comparing to control animals which persisted until the last measurement at 33 months of age (P<0.01, P<0.05, Fig. 2b).

**Figure 2.**
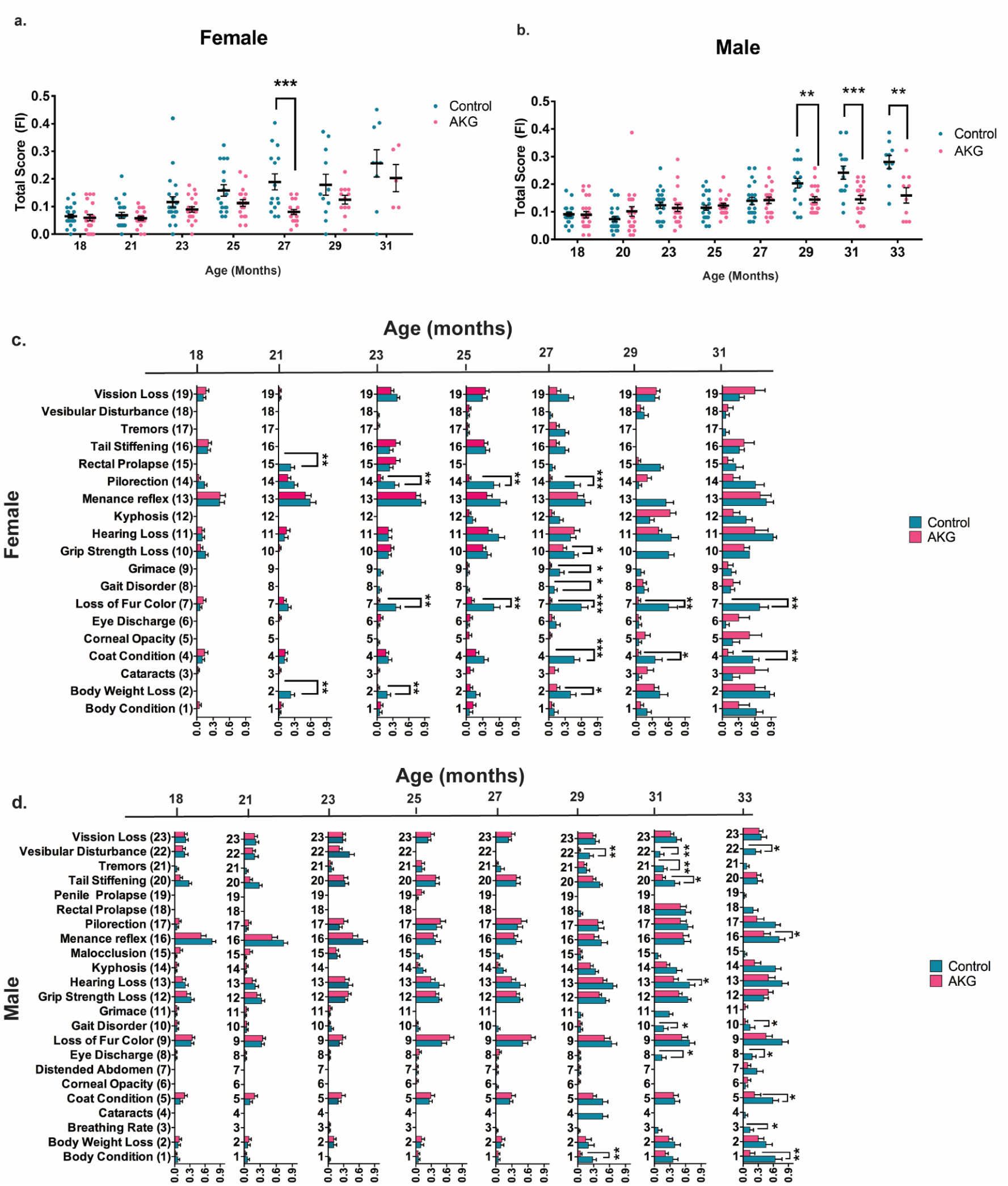
AKG treatment extends health span and alleviates age-associated frailty. Separately graphed (**a**) female and (**b**) male total FI scores during lifespan, comparing control mice (blue) to those fed AKG in the diet (pink) starting at 18th month. Each dot is the total score of one animal at specific age as indicated. Data are mean ±s.e.m. of each group. n= all animals alive at each measurement time, *P <0.05, **P< 0.01,***P< 0.001 (Two tailed t-test). Individually graphed frailty phenotypes that significantly change with age comparing control with AKG treated mice for (**c**) female and (**d**) male. Data are mean ±s.e.m. of the group, n= all animals alive at each measurement time. *P <0.05, **P < 0.01, ***P<0.001 (Two tailed t-test)

Since mice in our aging study were older than previously published papers on frailty [8], we first tested whether any individual frailty indicators show significant changes upon aging in our study. Our results show that aging significantly increased the incidence of 19 and 23 frailty phenotypes in female and male respectively (Two-way ANOVA, Extended Table 2). We reported the effects of CaAKG treatment on these phenotypes at different time points (Fig. 2c, d). CaAKG treatment significantly decreased the severity of multiple aging phenotypes in females including piloerection, grimace (pain assessment), loss of fur color, poor coat condition, grip strength loss, gait disorder, rectal prolapse and body weight loss (Fig. 2c). In males, grip strength loss, gait disorder, balance loss (vestibular disturbance), grimace, loss of menace reflex, hearing loss, eye discharge, poor coat condition, poor body condition, tail stiffening and abnormal breathing rate were all decreased (Fig. 2d). The improvement in health of both sexes was most prominent around the median life of the animal. Among the frailty measures affected by CaAKG, protection from age-related changes in female coat color was particularly prominent. CaAKG treatment could reverse age-dependent hair graying in first cohort, however in the second cohort of females, CaAKG treatment only prevented the hair graying (Extended Fig. 4). Gait improvement in CaAkG treated females and males close to median life is consistent with increased locomotion activity, as scored using metabolic cages (Extended Fig. 5). Interestingly, despite increased locomotion, the levels of oxygen consumption, carbon dioxide production and energy expenditure were significantly lower in the CaAKG treated group. The decrease in carbon dioxide production was preserved later in life (Extended Fig. 6). Advanced age-related weight loss has been reported in old human and rodents, which has been associated with morbidity [23]. Data from the second cohort of mice confirms that CaAKG administration leads to body weight preservation in male mice (P value<0.001, Extended Fig. 1b, d). Not all phenotypes were improved by CaAKG. For instance, treated mice failed to perform better in a treadmill exhaustion test and showed no cardiac functional improvement, as determined using echocardiography. Importantly, however, we did not detect any significant adverse changes associated with CaAKG treatment (Extended Fig. 7).

Since the age for onset of age-related phenotypes can be quite heterogeneous in mammals, we plotted frailty datasets not only as a function of chronological time, but also in proportion to the lifespan of each mouse by binning scores within ten percentile (e.g. scores collected between 60% and 70% of the animals’ lifespans were binned and plotted together (Fig 3). This allowed us to align assessments with respect to the biologic age of the animal. Our findings show that AKG treatment decreases the proportion of life in which the animal is frail and vulnerable to adverse health incomes (determined as the area under the frailty curve and calculated at a 52% reduction for females and 40% for males, Fig. 3 a, b, P<0.01). Our results also suggest that CaAKG modulates the rate of frailty changes with age (Fig. 3, Table). This improvement in healthy days of life is disproportionately larger than the increase in lifespan. Through the extensive frailty data collection, we have therefore been able to demonstrate that an intervention by a natural product results in properties consistent with morbidity compression.

**Figure 3.**
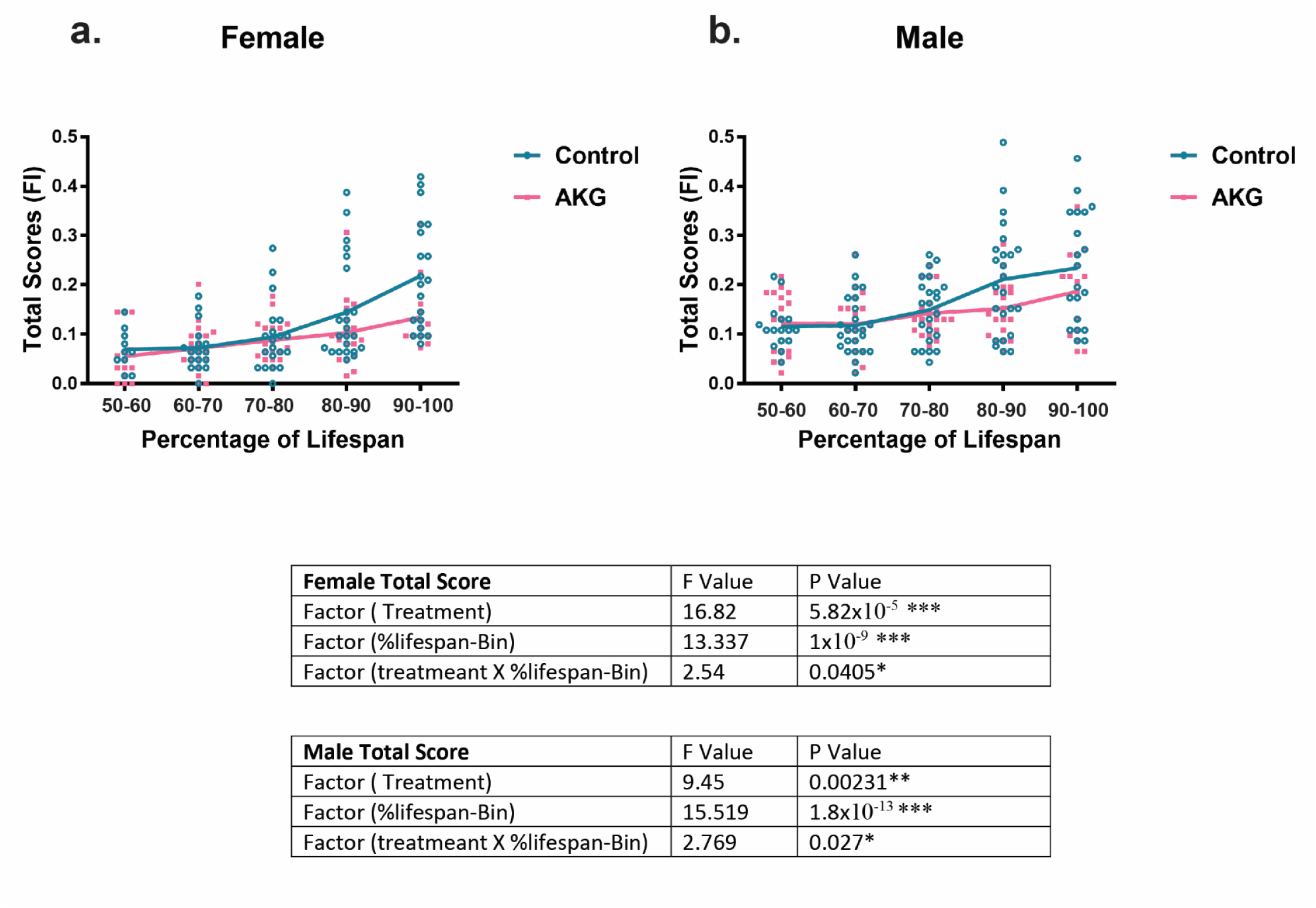
Compression of morbidity by AKG treatment. As the animal ages and gets closer to death (higher percentage of lifespan) it manifests several aging phenotypes and will be at its highest multi-morbidity risk, FI is Fraity Index and total scores of 31-phenotypes are considered as morbidity score. Separately graphed (**a**) female and (**b**) male mice total frailty scores as their percentage of lifespan. AKG treatment postpones the occurrence of aging phenotypes during lifespan and compresses the morbidity risk into fewer days of life in both sexes. Each dot is the total score of one animal. Lines are mean ±s.e.m. of the group. n= all animals alive at each time. Two-way ANOVA was used for comparison, The significant P value for treatment and (treatment × lifespan) suggest that AKG treatment modulates the rate of frailty changes with age.

To gain better mechanistic insight on the beneficial effects of AKG, we initiated AKG treatment in a new cohort of female mice at 18 months of age. The reported beneficial effect of AKG on lifespan in *C. elegans* was associated with reduced TOR signaling [12]. However, in mice we did not detect any decrease in mTORC1 signaling upon three months of AKG treatment (Extended Fig. 6). Age-associated diseases are accompanied by chronic inflammation, which is generally linked to an age-associated functional decline in mice and humans [24]. We measured levels of 31 inflammatory cytokines in the serum of aged female mice (28 months old). In untreated mice, the levels of most cytokines increase; however, CaAKG fed animals were largely refractory to these changes (Fig. 4a). There is a general trend of suppression for all the cytokines assayed in plasma of CaAKG treated animals (Fig. 4a, P<0.01). The reduction in inflammation is consistent with previous findings in the intestine of young pigs receiving AKG supplemented diet [25].

**Figure 4.**
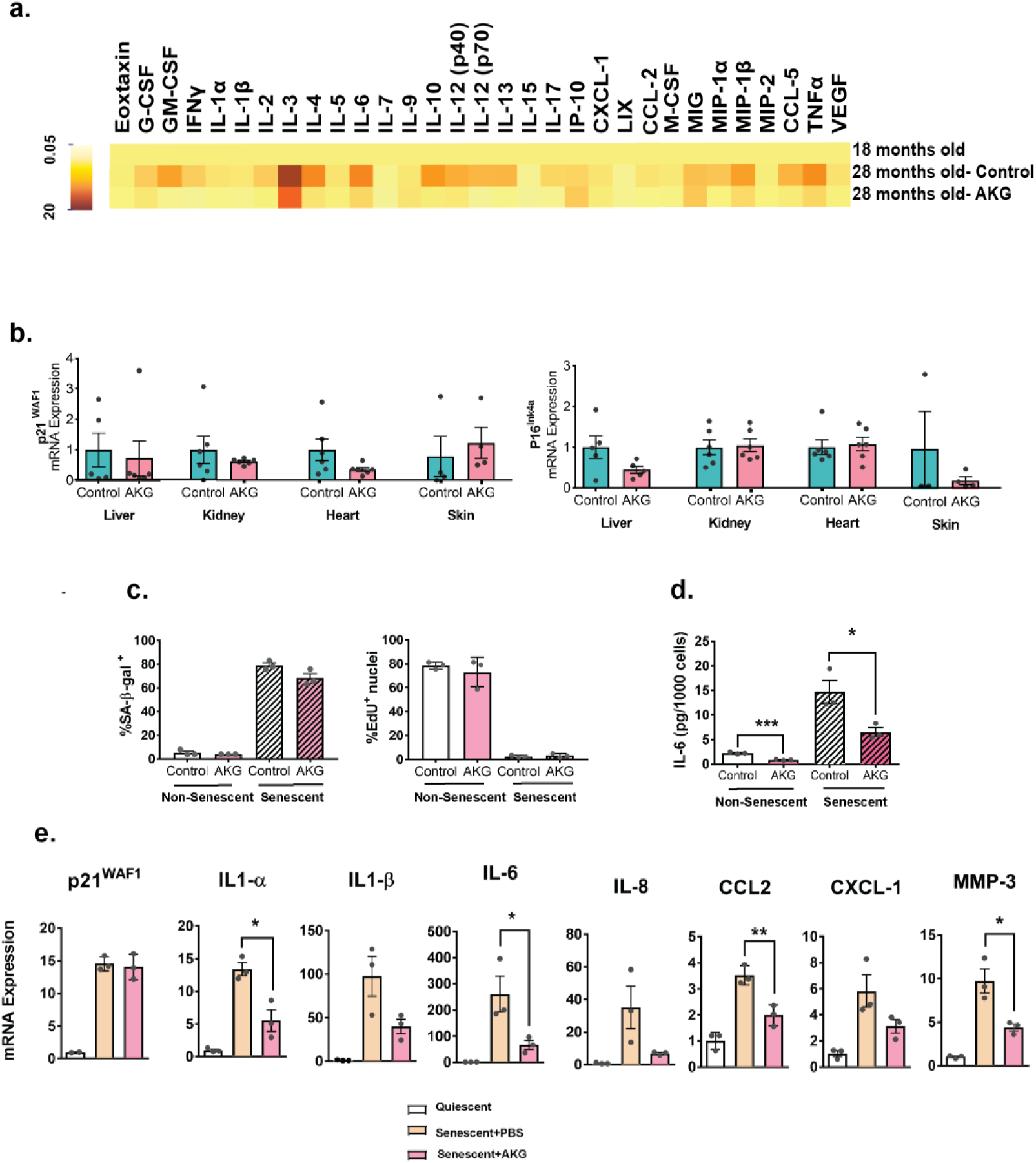
AKG reduces inflammation and lowers the proinflammatory SASP without preventing the senescence growth arrest. (**a**) Heat map of 31 inflammatory cytokines and chemokines from plasma of middle aged (female, age=18 months, n=11), aged control and AKG fed (female, 28 months old, n=5) animals. Cytokines show a general trend of reductions in AKG treated group comparing to aged control, for each treatment group, fold changes of all cytokines were calculated using the untreated 18 months old animals as reference. Values were all added together and was compared, **p =0.0046 (t-test two tailed). (**b**) qRT-PCR analysis of the indicated tissues of mice. (**c**-**e**) Ionizing radiation (IR) was used to induce senescence in IMR-90 fibroblasts. Cells were concurrently treated with PBS (control) or 1 mM AKG and were either mock (0 Gy) or irradiated (10 Gy). All assays were performed 10 days post irradiation. (**c**) Cells were stained for senescence-associated ß-galactosidase activity (left panel) or EdU incorporation (right panel). (**d**) IL-6 levels in conditioned media were determined by ELISA, normalized to cell number. (**e**) qRT-PCR analysis showing expression of senescence-associated secretory phenotype (SASP) genes, normalized to actin. Each dot is one independent experiment. Data are mean±s.e.m, *p < 0.05, **p<0.01 (t-test two tailed).

Studies have shown the accumulation of senescent cells in different tissues of old mice [26]. These cells can contribute to age-associated chronic inflammation by acquiring senescence-associated secretory phenotype (SASP) and their removal can extend mouse lifespan [27]. We looked into senescent markers and did not detect any significant changes for senescent markers (Fig. 4b). We performed cell culture studies to explore possible effects of NaAKG in primary fibroblasts. While no changes in the senescence formation response to ionizing radiation were observed; however, consistent to in-vivo findings we detect a significant reduction in a range of inflammatory cytokines, indicating that AKG can alter the SASP and reduce inflammation without effecting the formation of senescent cells (Extended Fig. 4c-e). Specifically, we found a reduction of IL-1b, IL-6, CCL2 and MMP3 without any changes in ß-gal and p21 (Fig. 4d, e).

## Discussion

Recent aging research has been based on the principle that there are underlying biological processes driving aging and that intervening in those processes will increase health and function, while delaying associated co-morbidities. Here we demonstrate that AKG, a key metabolite in TCA cycle, has longevity effects consistent with compressed morbidity in a long-term study of mouse aging. It is important that CaAKG administration started at 18 months of age still had robust effects. This is valuable since human clinical studies are likely to be initiated at a similar relative time point during aging. If translated to humans, this effect would be an ideal outcome, extending lifespan but more importantly reducing the debilitating period of functional decline and disease management that plague many aging individuals. Interestingly, in humans plasma AKG levels decline 10 folds between the ages of 40 and 80 [28]. The molecule is not available in the human diet, making direct supplementation the only feasible route to restore levels. AKG has been used in human clinical studies linked to diseases without associated adverse effects [29-31]. More studies will be needed to determine whether chronic AKG consumption can affect other parameters of aging. Given its GRAS status and human safety record, our findings point to a potential safe human intervention that may impact important elements of aging and improve quality of life in the elderly population.

## Material and methods

### Animal Housing and Diet

All mice were housed on a 12-h light/dark cycle and kept at 20–22 °C. Two independent cohorts of C57BL6/J mice were purchased from Jackson Laboratories at 14 months of age. Animals were aged on a regular mouse diet (Teklad Irradiated 18% protein and 6% fat diet-2918) which is very similar to their diet at Jax 5K52 (18-19% protein and 6-7% fat) prior to arrival, until they reached 18^th^ month of age when treatment started. AKG treated animals were subjected to a lifelong 2% (w/w) AKG supplement on 2918 diet while Control group were kept on standard-2918 diet. Pure calcium 2-oxoglutarate was purchased from Carbosynth Company and homogeneously mixed during manufacturing of the 2918 diet prior to irradiation and pelleting. Mice were housed in groups (5 per cage at a maximum) and aggressive male mice were isolated to prevent fighting. All lifespan and healthspan experiments were started around 18^th^ month of age. Mice were inspected daily, and treated for non-life threatening conditions as directed by the veterinary staff. The only conditions that received treatment were dermatitis and prolapse (topical solution three times per week). Total of 8 mice in control group and 3 mice in AKG group were treated for dermatitis and prolapse. The principal endpoint of the health span and lifespan study was natural death. The Buck Institute is a AAALAC accredited facility. Each room contains sentinel mice (CD-1 females-1 cage/75-100 cages). Health screening is done 4 times per year at 3 months intervals. Diagnostics consist of serological screening and fecal and fur analysis for internal and external parasites.

### Survival

The principle endpoint of health span and lifespan study was natural death. We recorded the age at which mice were found dead or selected for euthanasia (a procedure for mice deemed unlikely to survive for the next 48 hours and are in enormous discomfort). The criteria for euthanasia was based on an independent assessment by a veterinarian, according to AAALAC guidelines and only where the condition of the animal was considered [3]. Severe lethargy, rapid weight loss (over two weeks>20%), severe distended abdomen and body condition score with signs of pain (grimace), inability to move despite the stimuli, severe ulcer or bleeding tumor, severe temperature loss with abnormal breathing rate. Animals found dead or euthanized were necropsied for pathology score. No invasive measurements were performed on this population, n=180 animals (two cohorts of 90 animals). A sacrificed group were purchased at 14^th^ months of age, baselined and grouped the following week. The mice, n=12 were either receiving Teklad-2918 or 2% w/w AKG supplemented 2918. Animals were sacrificed and tissues were collected after 3 months of treatment. The Institutional Animal Care and Use Committee (IACUC) at the Buck Institute approved all animal experimental procedures, housing and diets for Research on Aging. Food intake and body weight were measured on a biweekly and bimonthly basis for the duration of the study.

### Baselining and grouping of the animals

Mammals age heterogeneously and the 18 months old mice already manifest some age-associated deterioration of health phenotypes. All the animals were scored before grouping and all the 31 scores were applied to assign animals into different groups. A balanced partitioning of mice was done: for any given mouse in any given group, there are similar mice in all other groups. This allows any outcome of the study to be more related to experiments or the treatment rather than the inherent property of a group.

### Aging Index (frailty Index)

One can find the complete protocol as published before by Whitehead et al., 2014. For the subjective properties of the assessments, all measurements were completely blinded. These assessments indicate age-associated deterioration of health and include evaluation of the animal musculoskeletal system, the vestibulocochlear/auditory systems, the ocular and nasal systems, the digestive system, the urogenital system, the respiratory system, signs of discomfort, body weight and body surface temperature. 0 is assigned if no sign of frailty is observed and the animal is healthy for that phenotype. A moderate phenotype and a severe phenotype will be scored 0.5 and 1 respectively. Loss of temperature and weight were scored using standard deviation for our study (Extended Table 2). We should note that the recessive *Cdh23* allele (*ahl*) in C57BL/6J strain, when homozygous, leads to increased susceptibility to age related hearing loss. Hearing loss was one of the most abundance aging phenotype in our data set.

### Statistical analyses

Python Software was used to extract all the healthspan data and create files compatible with R software for analysis. Data were analyzed using R, GraphPad Prism 7 and OASIS 2 software. Log-rank (Mantel–Cox) tests were used to analyze Kaplan–Meier curves, and a Fisher’s exact test was performed for maximum lifespan analysis (at 90% survival). Two-tailed Student’s *t*-tests were used for analyses of scoring at each time point between control and AKG treated group. Two-way analysis of variance (ANOVA) with Bonferroni post hoc correction was used for analysis between morbidity curves. The area under curve (AUC) of mortality graphs were measured baselining at 18 months of age. The changes in AUC were used to calculate the percent compression of morbidity.

### Inflammatory cytokines and chemokines

Blood were from the jugular vein of young (18 months old), aged control and AKG fed (29 months old) animals. The samples were sent to Eve technologies (Calgary, Alberta, Canada) for measurement of soluble cytokines and chemokines in serum using multiplex lase bead array technology (MD31).

### Cell culture

IMR-90 fetal lung fibroblasts were obtained from ATCC and were cultured at 37° C in 3%O_2_ and 5% CO^2^. Dulbecco’s modified Eagle’s media (DMEM) supplemented with 10% fetal bovine serum and streptomycin/penicillin were used. Media was changed every 2 days during the experiment. For damage-induced senescence, cells were irradiated with doses of either 0 or 10 Gy of ionizing radiation (IR). Cells were concurrently treated with PBS (control) or 1 mM AKG for 10 days, changing media every 2 days. All assays were performed 10 days post irradiation. EdU (5-ethynyl-2’-deoxyuridine) staining Proliferation Kit (iFluor 488) ab219801 was used to detect cell proliferation. Cells were stained for the senescence-associated β-gal (SA-β-gal) marker as described [3]. Non-senescent cells (having undergone fewer than 35 population doublings) were made quiescent by washing with PBS and incubating in DMEM containing 0.2% serum for 4 day. Cultures that had > 80% SA-β-gal positive cells and ≤ 4% EdU positive cells were considered senescent.

### ELISA

Conditioned media were prepared by washing cells 3 times in PBS and incubating them in serum-free DMEM containing penicillin/streptomycin for 24 h. Conditioned media were removed and cells were trypsinized for cell counts. The conditioned media were then centrifuged to remove cellular debris, and supernatants were used for ELISA. IL-6 ELISAs were performed using kits and procedures from R&D (#D06050). The resultant data were normalized to cell number.

### RT PCR

For cell culture experiments, RNA was isolated using ISOLATE II RNA mini kit (Bioline #BIO-52073). RNA quality and quantity were assessed using NanoDropTM 1000 Spectrophotometer measures (Thermo Scientific). Total cDNAs were synthesized from 500ng of RNA using random primers and iScript RT reagents following the manufacturer’s protocol Superscript II (Invitrogen, Carlsbad, USA). Gene expression was measured from cDNA using the Roche Universal Probe Library system (Indianapolis, IN, USA). All values were normalized to beta-actin.

For in-vivo study tissues were collected form 12 animals described as sacrificed group. Tissues were homogenized in 1 ml Invitrogen TRIzol™ Reagent using metal beads combined with high-speed shaking (Tissuelyser Qiagen at 20 Hrtz, for 6 min). Skin samples were crushed with pistol and liquid nitrogen prior to homogenizing step. The chloroform extraction and ethanol precipitation were performed on homogenized tissues to extract RNA. The RNA quality and quantity were assessed and cDNA were synthesized as described. Gene expression was quantified by real-time quantitative PCR using the Roche Universal Probe Library system (Indianapolis, IN, USA). The primer sets (0.1 µM) were as follows: 1) p16 F:5’-AACTCTTTCGGTCGTACCCC-3’ and R: 5’-TCCTCGCAGTTCGAATCTG -3’ with Custom designed probe : 5’-/56-FAM/AGG TGA TGA/ZEN/TGATGGGCAACGTTCAC/3IABkFQ -3’. 2) p 21 R: 5’-TTTGCTCCTGTGCGGAAC -3’ and F:5’-TTGCCAGCAGAATAAAAGGTG -3’ with probe #9. Transcript levels were normalized to Beta-glucuronidase (GUSB) as an endogenous control.

### Extended Data

Extended Data include eight figures and can be found here.

Unpublished data can be obtained by contacting the corresponding authors.

## Data availability

All other data are available from the corresponding authors on reasonable request.

## Acknowledgements

This work was supported by The Weldon Foundation and also by Ponce de Leon Health.

