## Supplemental Data for "Alpha-ketoglutarate, an endogenous metabolite, extends lifespan and compresses morbidity in aging mice"

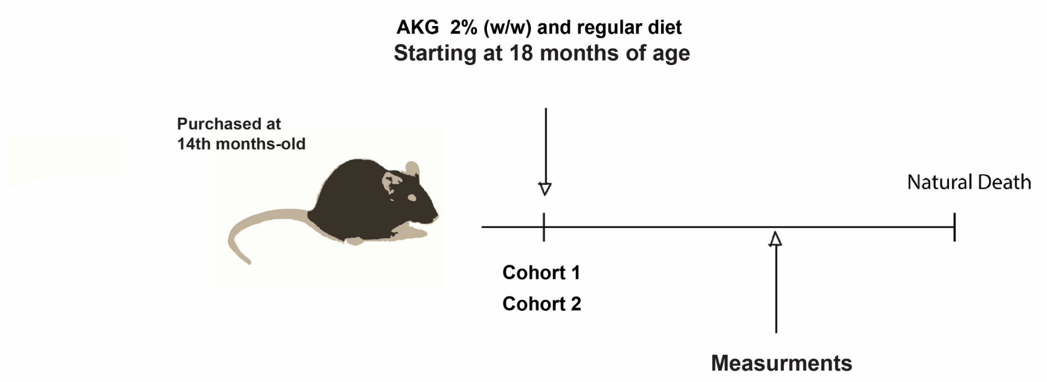


| | | |
|---|---|---|
| **Cohort-1** | n= 89 animals<br>(n= 42, female and n=46, male)<br>Age at start of the treatment:<br>18th Months old<br><br>Treatment Duration: Life long | • Measuring 31 Clinically relevant frailty index (FI)<br>• Lifespan |
| **Cohort-2** | n= 93 animals<br>(n= 45, female and n=48, male)<br>Age at start of the treatment:<br>18th Months old<br><br>Treatment Duration: Life long | • Lifespan<br>• Weight and body compositions<br>• Metabolic cages<br>• Heart Function |
| **Sacrificed group** | n= 12 (female)<br>Age at start of the treatment:<br>18th Months old<br>Treatment Duration: 3 months | • Blood collection at 28 months of age |

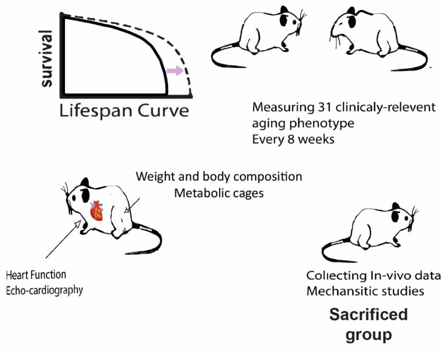


**Extended Data Diagram 1 | Experimental Design.** The mice were fed AKG or Standard diet at 18th months of age. The study consists of two cohorts ( n=182 animal total) and a sacrifice group (n=12). Cohort-1 mice were used for all the frailty index measurments and lifespan. Cohort-2 mice were started on diets at the same age for replication of survival, metabolic studies and complementary aging studies.

| | | Age (days) at 50% mortality | | Age (days) at 90% mortality | | Age (days) at 100% mortality | |
|---|---|---|---|---|---|---|---|
| | | **Cohort-1** | **Cohort-2** | **Cohort-1** | **Cohort-2** | **Cohort-1** | **Cohort-2** |
| **Female** | **Control** | 876 | 847 | 1015 | 931 | 1019 | 948 |
| | **AKG** | 932 | 875 | 1109 | 962 | 1116 | 975 |
| | **Percentage survival increase from inception of study (AKG vs. Cntrl)** | 16.6% | 10.5% | 19.7% | 8% | 21% | 6.6% |
| | **Percentage survival increase from birth (AKG vs. Cntrl)** | 6.3% | 3.3% | 9% | 3.3% | 10% | 2.8% |
| **Male** | **Control** | 955 | 812 | 1068 | 969 | 1109 | 987 |
| | **AKG** | 995 | 847 | 1155 | 987 | 1173 | 1008 |
| | **Percentage survival increase from inception of study (AKG vs. Cntrl)** | 9.6% | 12.8% | 16.4% | 4.2% | 11.2% | 4.7% |
| | **Percentage survival increase in survival from birth (AKG vs. Cntrl)** | 4.2% | 3.7% | 8.1% | 1.8% | 5.7% | 2.1% |

**Extended Data Table 1 | The effect of AKG treatment on lifespan**

| Integument: | Vestibulocochlear/Auditory: | Respiratory system: |
|---|---|---|
| 1. Alopecia<br>2. Loss of fur color<br>3. Dermatitis<br>4. Loss of whiskers<br>5. Coat condition | 14. Vestibular disturbance<br>15. Hearing loss<br>16. Cataracts<br>17. Eye discharge/swelling<br>18. Microphthalmia<br>19. Corneal opacity<br>20. Vision loss<br>21. Menace reflex<br>22. Nasal discharge | Breathing rate/depth |
| **Physical/Musculoskeletal:** | **Digestive/Urogenital:** | **Discomfort and others:** |
| 6. Tumors<br>7. Distended abdomen<br>8. Kyphosis<br>9. Tail stiffening<br>10. Gait disorders<br>11. Tremor<br>12. Forelimb grip strength<br>13. Body condition score | 23. Malocclusions<br>24. Rectal prolapse<br>25. Vaginal/uterine/ Penile prolapse<br>26. Diarrhea | 28. Mouse Grimace<br>29. Piloerection<br>30. temperature*<br>31. Weight* |

**Extended Data Table 2 | List of assessed deficits in aged mice- Frailty index.** All the phenotypes were scored as described before by Whitehead, J.C., et al., except for temperature and weight. ***Temperature and Weights Scoring:** Briefly, for weight and temperature, the new scaling scores were used; Average and standard deviations (STDEV) were calculated sex specifically using our own baseline data sets (data collected before the start of the treatment-mice at 18th month age). A decrease in temperature [1] or weight ( Extended figure 1b, d and [2]) within one STDEV is scored as a (0), a decrease bigger than one STDEV but smaller than 2 STDEV will be scored as a (0.5) and any decrease more than 2 STDEV is scored as a (1)

Table 3

| Frailty Phenotypes (Female) | Change with Age | | Frailty Phenotype (Female) | Change with Age | |
|---|---|---|---|---|---|
| | F-Value | P-Value | | F-Value | P-Value |
| **1. Alopecia** | 1.614 | 0.144424 | **17. Kyphosis** | 14.393 | $8.58 \times 10^{-14}$ *** |
| **2. Body Condition** | 10.585 | $2.68 \times 10^{-10}$ *** | **18. Loss of Whiskers** | 0.244 | 0.961 |
| **3. Body Weight** | 10.099 | $8.36 \times 10^{-10}$ *** | **19. Malocclusion** | 1.632 | 0.139 |
| **4. Breathing Rate** | 1.631 | 0.1396 | **20. Menace Reflex** | 4.940 | $7.66 \times 10^{-5}$ *** |
| **5. Cataracts** | 8.571 | $2.3 \times 10^{-8}$ *** | **21. Microphthalmia** | 1.591 | 0.151 |
| **6. Coat Condition** | 2.818 | 0.011639 * | **22. Nasal Discharge** | 0.20 | 0.98 |
| **7. Corneal Opacity** | 9.491 | $2.96 \times 10^{-9}$ *** | **23. Piloerection** | 3.131 | 0.00579 ** |
| **8. Dermatitis** | 1.855 | 0.0898 | **24. Rectal Prolapse** | 5.236 | $4.68 \times 10^{-5}$ *** |
| **9. Diarrhea** | 2.255 | 0.13462 | **25. Tail Stiffening** | 9.467 | $3.13 \times 10^{-9}$ *** |
| **10. Distended Abdomen** | 1.972 | 0.0709 | **26. Tremors** | 2.167 | 0.0474 * |
| **11. Eye Discharge** | 2.703 | 0.015* | **27. Tumors** | 1.069 | 0.382 |
| **12. Loss of Fur color** | 4.744 | 0.000146 *** | **28. Temperature Loss** | 2.266 | 0.0717 |
| **13. Gait Disorder** | 6.495 | $2.57 \times 10^{-6}$ *** | **29. Vaginal Prolapse** | NA | NA |
| **14. Grimace** | 2.862 | 0.010541 * | **30. Vestibular Disturbance** | 3.633 | 0.00187 ** |
| **15. Grip Strength Loss** | 19.422 | $<2 \times 10^{-16}$ *** | **31. Vision Loss** | 9.458 | $3.22 \times 10^{-9}$ *** |
| **16. Hearing Loss** | 17.053 | $4.31 \times 10^{-16}$ *** | | | |

Table 4

| Frailty Phenotypes (Male) | Change with Age | | Frailty Phenotype (Male) | Change with Age | |
|---|---|---|---|---|---|
| | F-Value | P-Value | | F-Value | P-Value |
| **1. Alopecia** | 0.884 | 0.520 | **17. Kyphosis** | 22.588 | 2e-16 *** |
| **2. Body Condition** | 16.581 | < 2e-16 *** | **18. Loss of Whiskers** | 1.312 | 0.244 |
| **3. Body Weight** | 6.103 | 1.19e-06 *** | **19. Malocclusion** | 4.022 | 0.000314 *** |
| **4. Breathing Rate** | 2.341 | 0.0243 * | **20. Menace Reflex** | 3.394 | 0.00166 ** |
| **5. Cataracts** | 1.489 | 0.1705 | **21. Microphthalmia** | 2.431 | 0.01948 * |
| **6. Coat Condition** | .755 | 0.00868 ** | **22. Nasal Discharge** | 0.838 | 0.556 |
| **7. Corneal Opacity** | 3.116 | 0.00344 ** | **23. Piloerection** | 7.684 | 1.49e-08 *** |
| **8. Dermatitis** | 1.607 | 0.13280 | **24. Penile Prolapse** | 3.665 | 0.000814 *** |
| **9. Diarrhea** | NA | NA | **25. Rectal Prolapse** | 2.234 | 0.0315 * |
| **10. Distended Abdomen** | 9.313 | 1.86e-10 *** | **26. Tail Stiffening** | 3.118 | 0.00341 ** |
| **11. Eye Discharge** | 2.671 | 0.010713 * | **27. Tremors** | 2.832 | 0.00712 ** |
| **12. Loss of Fur color** | 5.881 | 2.02e-06 *** | **28. Tumors** | 0.707 | 0.66609 |
| **13. Gait Disorder** | 2.049 | 0.04892 * | **29. Temperature Loss** | 1.096 | 0.365 |
| **14. Grimace** | 8.456 | 1.85e-09 *** | **30. Vestibular Disturbance** | 7.562 | 2.11e-08 *** |
| **15. Grip Strength Loss** | 4.287 | 0.000154 *** | **31. Vision** | 2.943 | 0.00537 ** |
| **16. Hearing Loss** | 5.671 | 3.59e-06 *** | | | |

**Extended Data Table 3 and 4** | **List of all frailty phenotypes and their interaction with aging (time)**. P value is the result of two-way ANOVA for changes in frailty as a function of time for females (Table 3) and males (Table 4), *P <0.05, **P <0.01 and ***P <0.001.

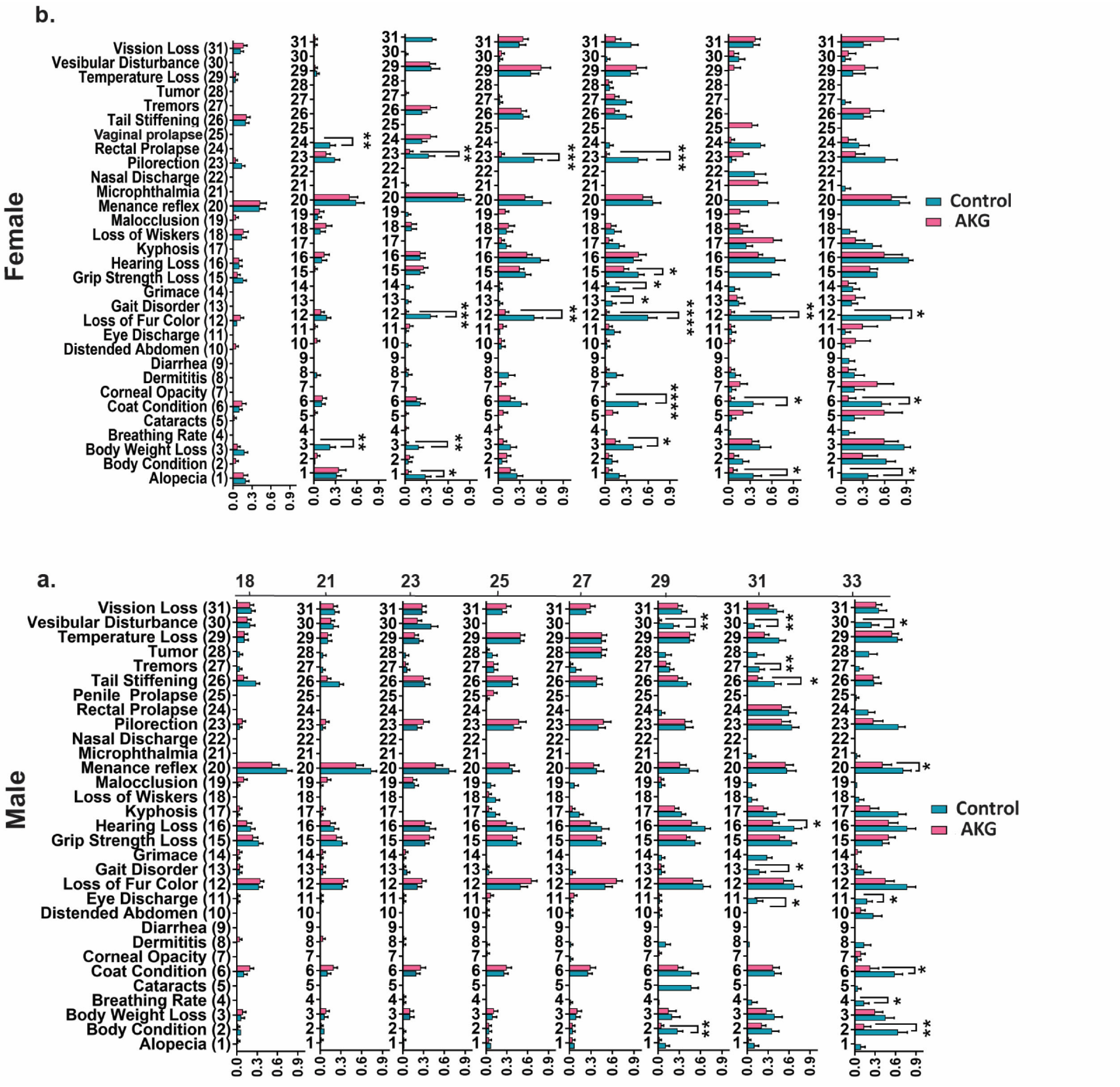


**Extended Data Figure 1** | **AKG treatment extends health span and alleviates age-associated frailty**. All individual frailty phenotypes (total of 31 phenotypes), separately graphed (**a**) female and (**b**) male. Data are mean ±s.e.m. of the group, n= all animals alive at each measurement time. *P <0.05, **P < 0.01, ***P<0.001, ****P <0.0001 (Two tailed t-test ).

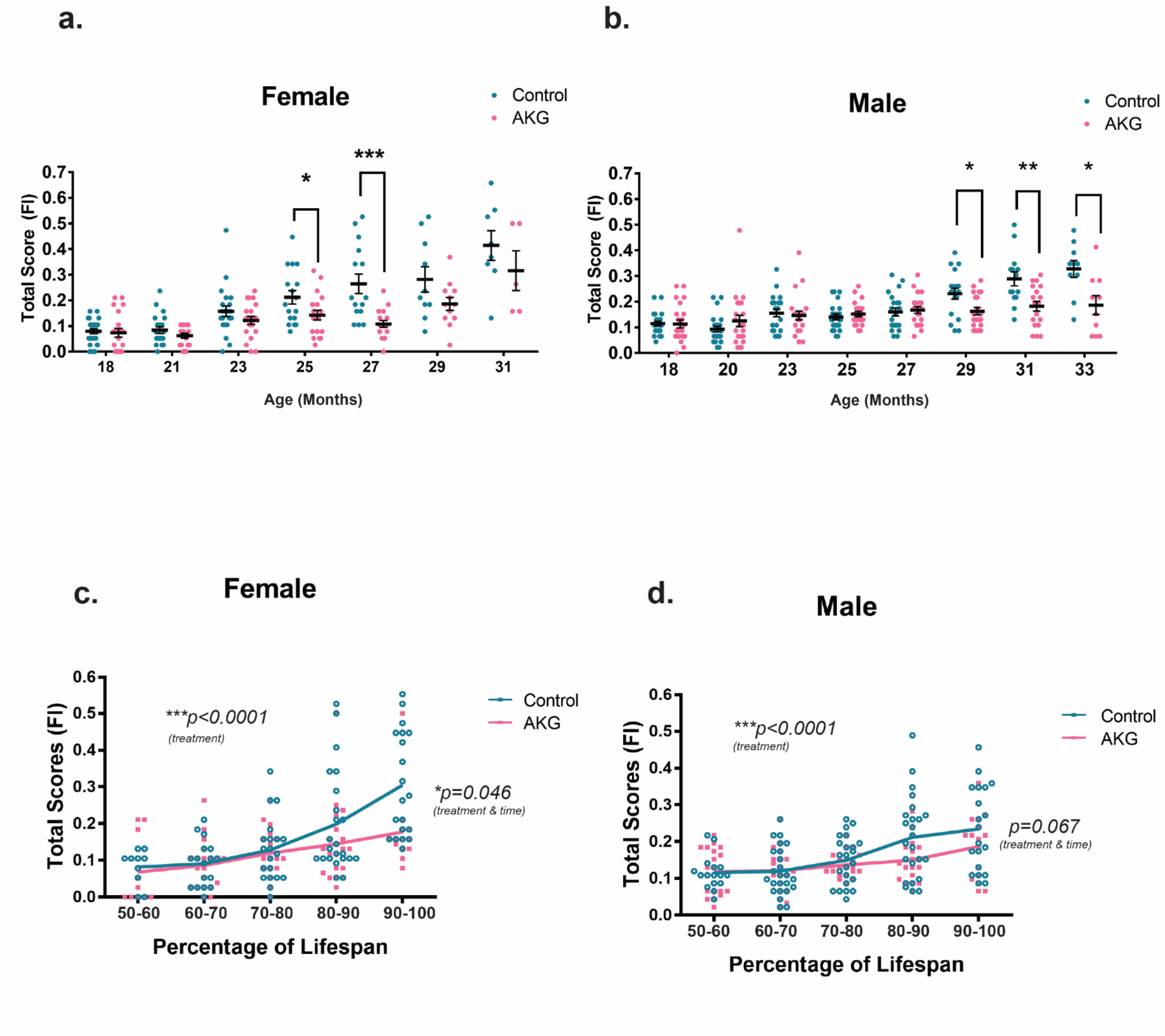


**Extended Data Figure 2 | Compression of morbidity by AKG treatment (including only the phenotypes significantly change with age).** Separately graphed (**a**) female and (**b**) male total FI scores. Each dot is the total score of one animal at specific age as indicated. The total score includes only the frailty phenotypes that significantly change with aging during lifespan (total of 19 phenotypes for female and 23 phenotypes for male). Data are mean ±s.e.m. of each group. n= all animals alive at each measurement time, *P <0.05, **P< 0.01 (Two tailed t-test). Separately graphed (**c**) female and (**d**) male mice total frailty scores as their percentage of lifespan. As the animal ages and gets closer to death (higher percentage of lifespan) it manifests several aging phenotypes and will be at its highest multi-morbidity risk, FI is Fraity Index and total scores are considered as morbidity score. AKG treatment postpones the occurrence of aging phenotypes during lifespan and compresses the morbidity risk into fewer days of life in both sexes. Each dot is the total score of one animal. Lines are mean ±s.e.m. of the group. n= all animals alive at each time.The morbidity (total score) show strong correlation with aging (***P < 0.001) and AKG treatment group show improvement in health indicted by total score (***P < 0.001). In addition, AKG treatment modulates the slope of morbidity by time in female (***P = 0.046). Two-way ANOVA was used for comparission, ***P < 0.001, *P<0.05.

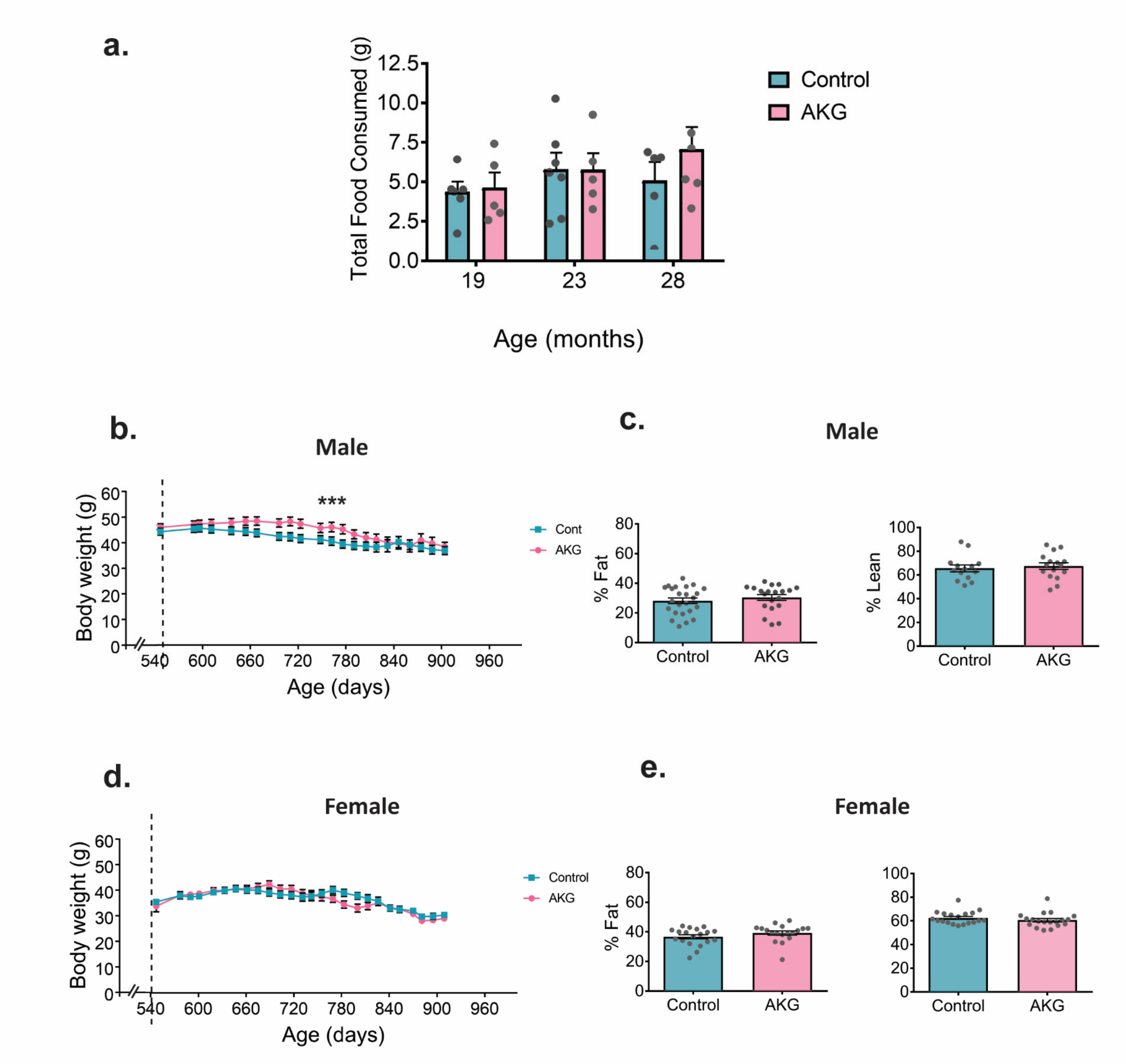


**Extended Data Figure 3 | AKG preserves body weight in male animals ( Cohort-2 data).** (**a**) Food consumption was measured for 3 consecutive days in different times during lifespan of control (n=6) and AKG fed (n=5) mice, the arrows at 19, 23 and 28 months show the age at which food consumption was measured. Data are mean±s.e.m, no significance change (t-test two tailed). (**b**, **d**) Longitudinal male and female body weights, n = all live animals in the study at each time points, data are mean±s.e.m. Two way ANOVA tests check if independent variables including time and treatment (AKG) affect male and female body weight. The comparison provided evidence of significant effects of both treatment and time on (**b**) male body weight. ***p <0.0001 and ***p <0.001 and effect of (**d**) time on female body weight ***p <0.001. (**c**, **e**) Male and Female body composition, n = all live animals in the study, data are mean±s.e.m

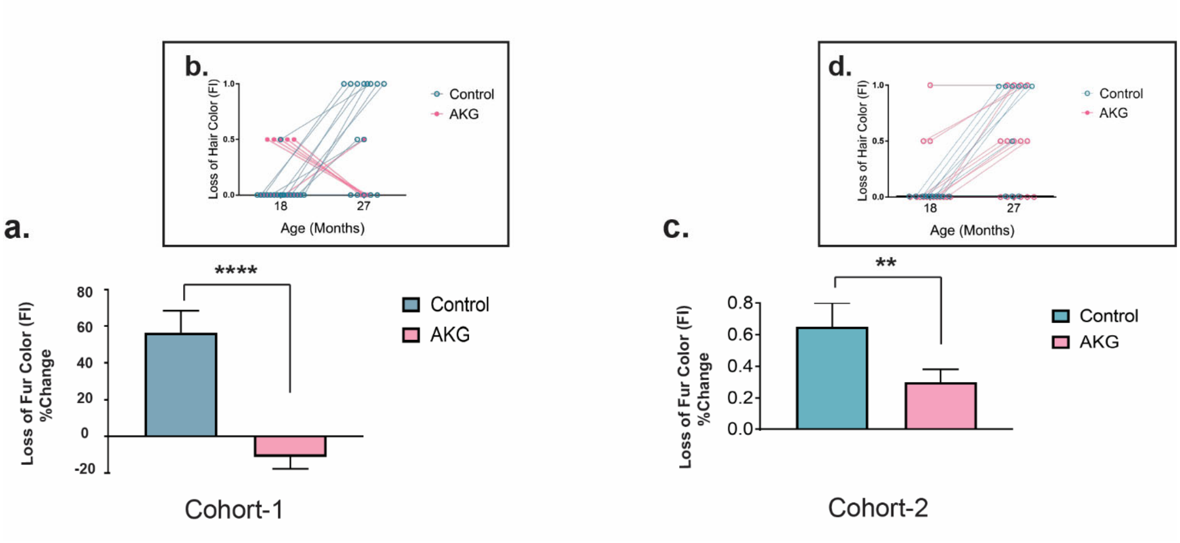


**Extended Data Figure 4 | AKG treatment strongly prevents age-associated hair discoloration in female mice. (a, c**) Overall assessment of changes in fur color index of mice in each treatment group after 9 months of treatment. Control (n= 15) and AKG (n=18), Data are mean±s.e.m. ****P<0.0001 and **P <0.01 (t-test, two tailed). (**b, d**) Each dot is a single mouse and connecting lines tie the base line score for each mouse to its scoring after 9 months of treatment. Aging caused an increase in gray hair in control mice. AKG grouped mice had more gray hairs at baseline ($18^{th}$ months of age) and AKG treatment could reverse the hair discoloration in first cohort and prevent the hair discoloration in the second cohort of mice.

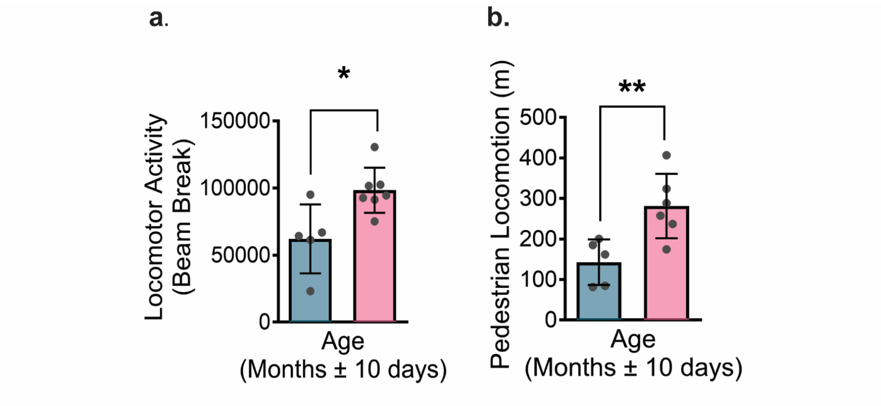


**Extended Data Figure 5 | AKG treatment improves locomotion in aged mice (Cohort-2 data).** (**a**, **b**) Locomotor activity and pedestrian locomotion was measured at the median life of the animals (28 months old). Female mice control (n=5) and AKG (n=6). Data are mean±s.e.m, *P value=0.014, **P value=0.00097 (t-test two tailed).

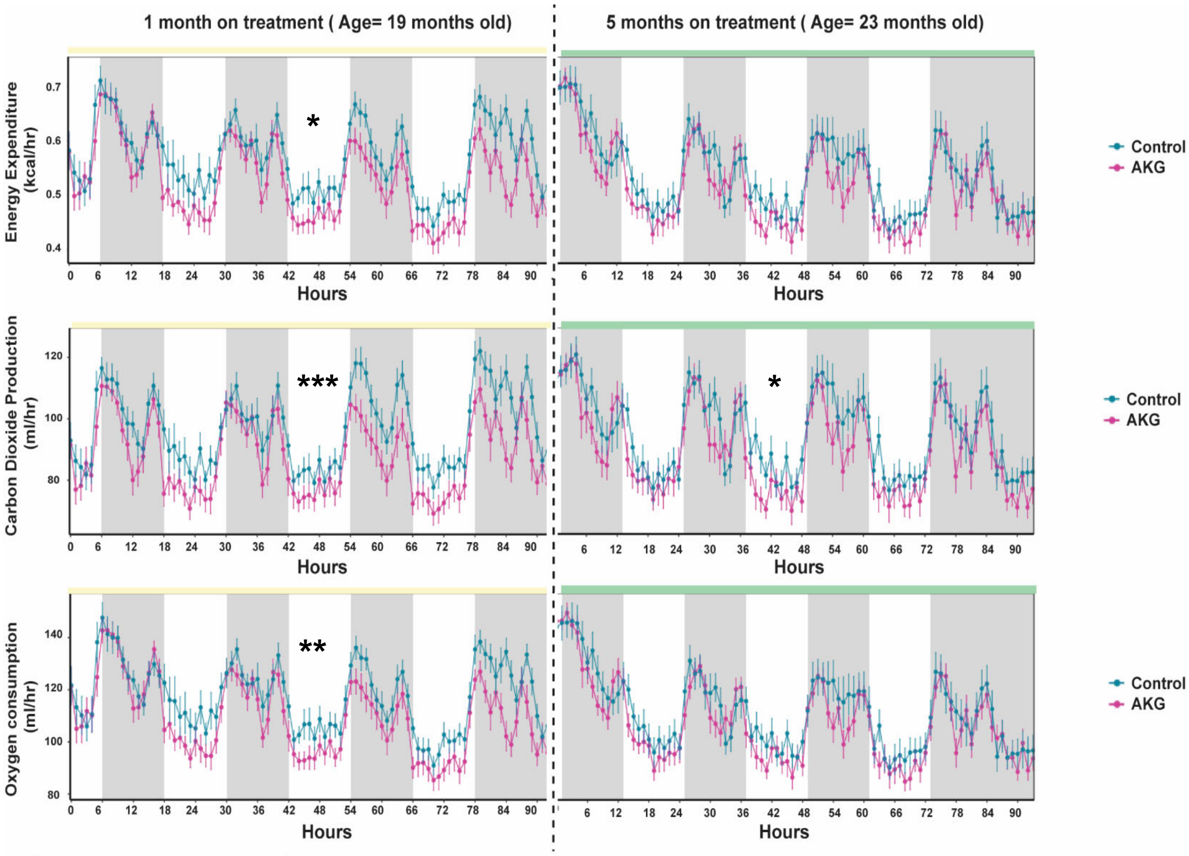


| P value | 1 month on treatment (19 months old) | | | 5 months on treatment (23 months old) | | |
|---|---|---|---|---|---|---|
| Energy Expenditure | Light = 0.11 | *Dark =0.021 | *Full Day= 0.0121 | Light = 0.12 | Dark= 0.304 | Full Day=0.058 |
| Carbon Dioxide | *Light =0.014 | *Dark=0.011 | *** Full Day<0.001 | *Light=0.04 | *Dark=0.037 | *Full Day=0.019 |
| Oxygen Consumption | *Light=0.045 | *Dark=0.014 | **Full Day= 0.006 | Light =0.16 | Dark=0.38 | Full Day= 0.25 |

**Extended Data Figure 6 | AKGdecreases the metabolic rate of aged mice (Cohort-2 data).** Oxygen consumption, Carbon dioxide production and Energy expenditure decrease upon AKG administration. Mice were monitored for about four consecutive days (92 hrs).The measurements were done at two separate time points during lifespan (19 and 23 months old) using same animals for both runs. Plots were generated using CalR, the data is adjusted to bodyweight. Control (n=5) and AKG (n=5). Data are mean±s.e.m. *$p < 0.05$ and ***$p < 0.001$ (Two way ANOVA tests).

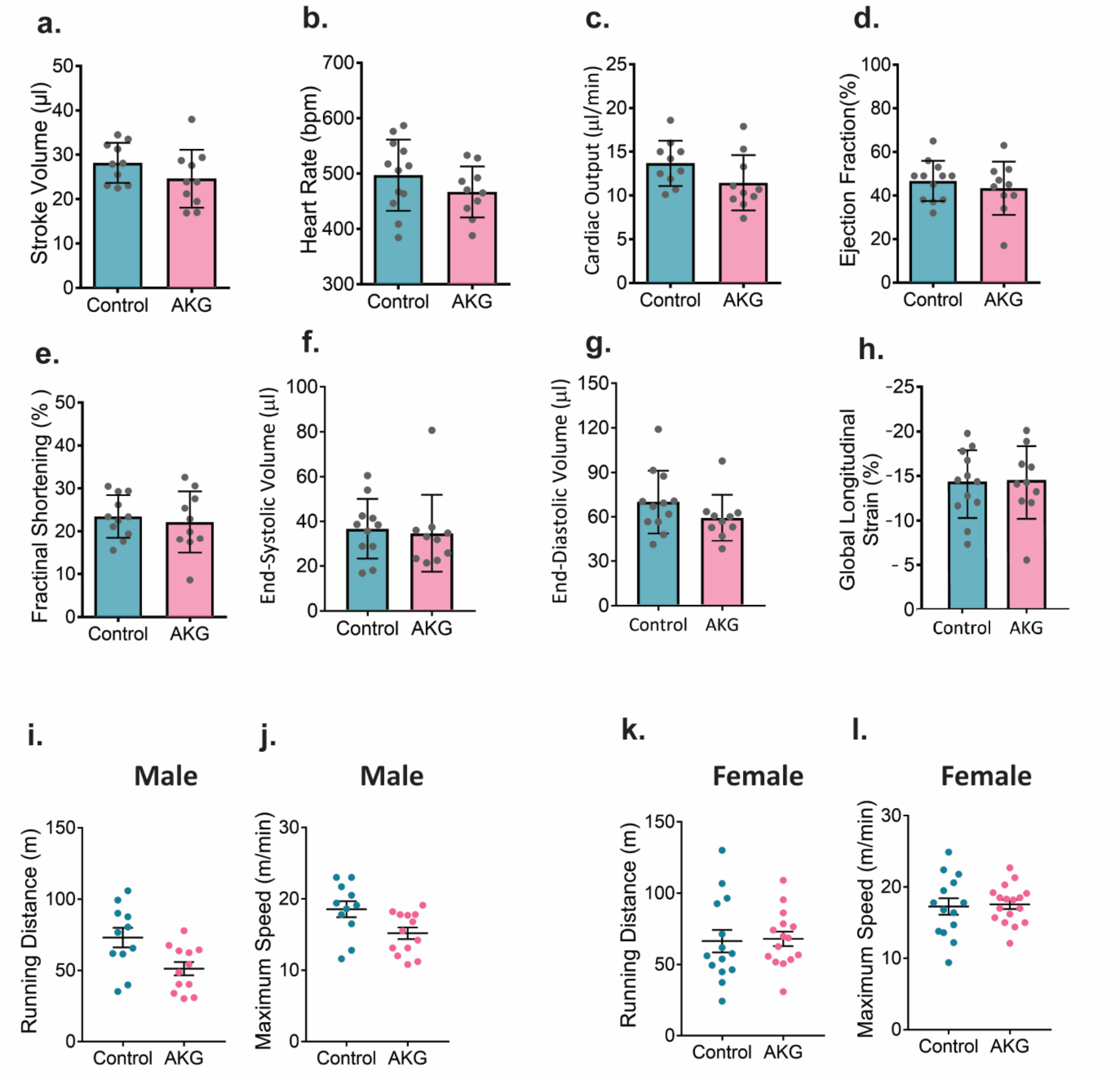


**Extended Data Figure 7 | AKG treatment does not improve heart function.** (**a-h**) Echocardigraphy test was performed to measure cardiovascular function close to animal median life, age=29 months old, n= all female animals alive at the time of study, data are mean±s.e.m. No significant change was observed for any of the measurements (t-test two tailed). Treadmill exhaustion tests were performed to measure cardiovascular system and motor function for (**i**,**j**) male and (**k**,**l**) female, age= 29 month old. n= all animals alive at the time of study. No significant change (t-test two tailed).

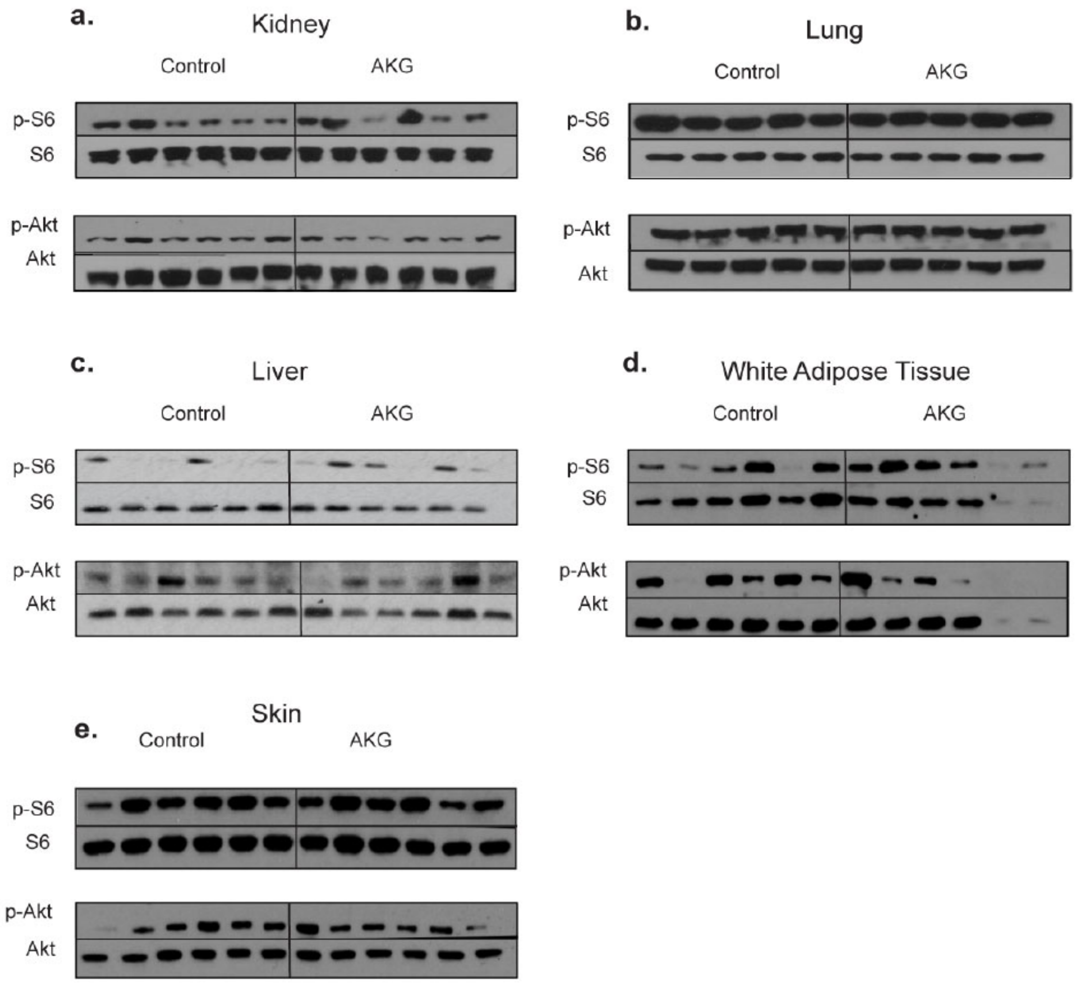


**Extended Data Figure 8 | mTOR activity is not changed in tissues of mice treated with AKG.** (**a-f**) Western blots of mTORC1 (indicated by p-S6/S6) and mTORC2 (indicated by p-Akt/Akt) activities in multiple tissues of mice treated with AKG for 3 months. Overall three months AKG treatment does not change mTORC1 and mTORC2 activities

**Material and Methods**

**Metabolic data**

Metabolism was measured applying indirect Calorimetry. We used Promethion metabolic cage system-Sable Systems International. The system is equipped with GA-3 small mammal gas analyzers for measurements of $O_2$ (consumption) and $CO_2$ (production). Energy expenditure, food intake, water consumption, body weight, physical activity and volunteer exercise were simultaneously recorded over 4 consecutive days (96 hours). Mice were housed individually in metabolic cages and accustomed to their environment a day before the start of recording. Data were analyzed using Sable System Expedata-P Data Analysis Software. Subsequently, we applied CalR software, a free web tool for analysis of experiments using indirect calorimetry [3], to analyze our raw data, generate some of our plots and run statistical analysis (https://calrapp.org). The whole-body composition analysis was conducted using a quantitative nuclear magnetic resonance machine (EchoMRI-2012, Echo Medical Systems).

**Transthoracic echocardiography**

Transthoracic echocardiography Echocardiography examination was performed using a high resolution (32-55 MHz) Visualsonics Vevo 2100 micro-ultrasound system with the echocardiography probe (MS-400). Individual mouse was placed on a heating pad (37°C) and minor sedation (0.5% isoflurane oxygen) was used to paralyze the animal (minimizing the cardiac suppression side effect) during the measurement time. Doppler imaging, 2D and M-mode echocardiography was performed to evaluate cardiac morphometry, systolic function, and Mean baseline myocardial performance index (MPI).

**Treadmill exhaustion test**

We modified the protocol by Beatriz castro et al., 2017 [4]. Since, mice were old ($28^{th}$ month-old) the initial speed and the acceleration were adjusted for our study. All mice were trained and adapted to the environment for three consecutive days for 10 min at 5m/min before the actual experiment. On the day of experiment mice were warmed up for 3 min at 5 m/min after which the speed was accelerated by 1.5 $m/min^{-2}$. We used air puff as stimuli to keep the animal running. The maximal speed and distance were recorded once the mice were exhausted (signs of exhaustion including heavy breathing, hunched back and unwillingness of the animal to get on the treadmill belt despite of 10 air puffs).

**Western**

Mouse were fasted overnight, the next morning tissues; heart, lung, kidney, adipose tissue (visceral fat in females, carefully along the epididymis and the uterus) and skin (back skin, on the spine midway between the head and the tail) were dissected and immediately frozen in liquid nitrogen. Tissues were homogenized using the Omni TH homogenizer (Omni International) on ice in Radioimmunoprecipitation assay (RIPA) buffer; 300 mM NaCl, 1.0% NP-40, 0.5% sodium deoxycholate, 0.1% SDS, 50 mM Tris (pH 8.0), protease inhibitor cocktail (Roche) and phosphatase inhibitor 2, 3 (Sigma). Samples were centrifuged at 13,200 rpm for 10 min, 4°C. The protein contents of the supernatants were assessed using the detergent compatible (DC) protein assay (Bio-Rad). Equal amounts of protein were resolved by SDS-PAGE (4%– 12% Bis-Tris gradient gel, Invitrogen), transferred to nitrocellulose membranes, and incubated with protein/phosphoprotein-specific antibodies. The antibodies against phosphorylated rsS6$^{S240/244}$ (5364), Akt$^{S473}$(4058), S6 (2217), Akt (4691), and glyceraldehyde 3-phosphate dehydrogenase (GAPDH; 2118) were purchased from Cell Signaling Technology. Protein bands were revealed using the Amersham enhanced chemiluminescence (ECL) detection system (GE Healthcare) and quantified by ImageJ software.


1. Talan, M., *Body temperature of C57BL/6J mice with age.* Exp Gerontol, 1984. **19**(1): p. 25-9.
2. Martin-Montalvo, A., et al., *Metformin improves healthspan and lifespan in mice.* Nat Commun, 2013. **4**: p. 2192.
3. Mina, A.I., et al., *CalR: A Web-Based Analysis Tool for Indirect Calorimetry Experiments.* Cell Metab, 2018. **28**(4): p. 656-666 e1.
4. Yorke, A., et al., *Development of a Rat Clinical Frailty Index.* J Gerontol A Biol Sci Med Sci, 2017. **72**(7): p. 897-903.
